# Spatial image gradient estimation from the diffusion MRI profile

**DOI:** 10.1101/2025.06.06.658348

**Authors:** Iman Aganj, Thorsten Feiweier, John E. Kirsch, Bruce R. Fischl, Andre J. van der Kouwe

## Abstract

**Purpose:** In the course of diffusion, water molecules experience varying values for the relaxation-time properties of the underlying tissue. This factor, which has rarely been accounted for in diffusion MRI (dMRI), is modeled in this work, allowing for the estimation of the gradient of the relaxation-time properties from the dMRI signal.

**Methods:** With the aim of mining the dMRI data for information about the spatial variations in the tissue relaxation-time properties, a new mathematical relationship between the diffusion signal and the spatial gradient of the image is derived, enabling the estimation of the latter from the former. The hypothesis was validated on human brain dMRI images from three datasets: the public Human Connectome Project Young Adults database, 10 healthy volunteers and 1 *ex vivo* sample scanned in-house with stimulated-echo diffusion encoding and a long diffusion time of 1 second (which will be made publicly available), and 3 subjects from the public Multi-TE database. The effects of the confounding factor of “fiber continuity” were furthermore measured.

**Results:** The image spatial gradient estimated from the diffusion signal was compared to the gold-standard spatial gradient approximated through the finite difference. The former gradient was significantly related to the latter in all datasets (i.e., with a difference significantly smaller than chance), with an effect distinct from fiber continuity.

**Conclusion:** The results support the hypothesized relationship between within-voxel dMRI signal and image gradient, with an effect that was not explainable by the confounding factor of fiber continuity.

## 1. Introduction

As a noninvasive imaging modality, diffusion-weighted magnetic resonance imaging (dMRI) provides a wealth of information that has been proven valuable in revealing the microarchitecture of the neural tissue (1,2). Many imaging biomarkers for neurodegenerative diseases have been derived from dMRI in the past few decades (3), notably through the *in vivo* quantification of structural brain connectivity (4,5). Developing mathematical models consistent with dMRI physics and suitable to analyze dMRI data, which are inherently high-dimensional, has been necessary to extract the desired information about the tissue from the images (6-8).

Per the standard MRI model described by the Bloch equations, the MRI signal is proportional to the mean proton density (PD) inside a voxel, weighted according to relaxation times (RTs) of the tissue (9). In dMRI, where additional diffusion-encoding gradients are imposed, the signal is further attenuated with the displacement of water molecules in the direction of these gradients. Such a change in the measured signal enables inference of the diffusivity of water along different orientations, and hence the estimation of tissue properties such as fiber orientations (10,11). While the tissue RTs and diffusivity are distinct properties, they have been shown to be related to each other both theoretically (12-15) and empirically (16-20). The relationship between the two has been particularly explored in the context of diffusion-relaxation correlation (21). A number of studies have also examined how the dMRI signal is confounded by the presence of local magnetic field gradients, especially by the effects of susceptibility, at the mesoscopic scale (22-26).

Given the finite resolution of MRI, i.e., the measured signal being the average of continuous values inside a voxel, inferring within-voxel variations of an estimated quantity is often nontrivial at best. It is thus common to make approximations in MRI signal modeling by discounting such within-voxel variations. An example of such a scenario, which is the focus of this work, is in dMRI signal modeling. Although the interplay between the diffusion of spin-carrying molecules and the measured RTs has been thoroughly discussed (12-15), the effects of the spatial variations of the latter have rarely been accounted for in dMRI modeling. An exception is the work by Novikov *et al* (15), where the effects of the intrinsic variations of the transverse relaxation rate on the measured dMRI signal have been rigorously examined, and a general integral relationship between the apparent diffusion coefficient (ADC) and the Fourier transform of the autocorrelation function of the transverse relaxation rate is provided. Besides that, standard dMRI models assume the RT properties of the tissue (resulting in T_1_ and T_2_ weighting) to be constant inside the voxel, an assumption that disregards the (small) contribution of the within-voxel variations of these properties to the dMRI signal.

In this work, we formulate and test a hypothesis based on the premise that, in the course of diffusion, water molecules experience varying values for the RT properties of the underlying tissue. We hypothesize that the spatial variation in tissue RTs (which is related to image gradient) affects the dMRI signal, thereby creating a relationship between the diffusion profile measured by dMRI and the spatial gradient of the image intensity. We derive this mathematical relationship and propose an approach to estimating the spatial gradient of the image from the dMRI signal (independently at each voxel). We validate our model via experiments on human brain dMRI data, specifically public images from the Human Connectome Project (HCP) Young Adult database (27), stimulated-echo (STE) (28,29) images that we acquired at our Center (see Section 6 for public availability of our data), and the public Multi-TE (MTE) dataset (30). The STE data were acquired with a very long diffusion time of one second, which our model predicts should increase the hypothesized effect size.

We have previously presented a preliminary abstract of this work (31). In the following, we will describe our model and methods (Section 2), present experimental results (Section 3), discuss them (Section 4), and provide our code and data (Section 6).

## 2. Methods

### 2.1. Signal Modeling

The Stejskal-Tanner pulsed gradient spin-echo sequence (10) applies two gradient pulses 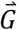 of duration δ, separated in time by Δ. Molecules located at 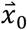 during the first pulse and ending up at 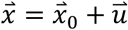 at the second pulse presumably contribute the following to the dMRI signal 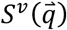 at voxel *v*, where 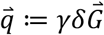 is the *q*-vector (or *2π*times it with an alternative definition) with *y* the gyromagnetic ratio (32):

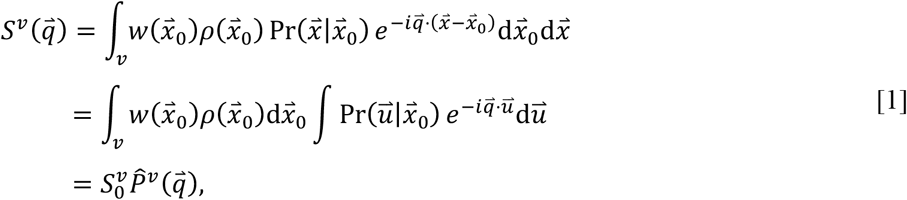

where *ρ* is PD, and 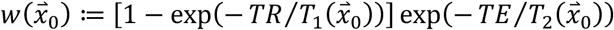 is the RT weighting for the repetition time *TR*, echo time *TE*, and the tissue longitudinal and transverse RTs, 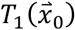 and 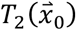, respectively. 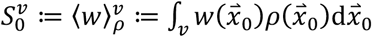 is the baseline non-diffusion-weighted (i.e., b=0) image, where 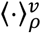 denotes *ρ*-weighted sum inside *v*, i.e. the voxel. 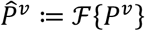 is the Fourier transform of 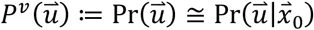, which is the probability of diffusion with the amount 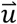 (a.k.a. ensemble average propagator) during the effective diffusion time *τ* := Δ − *δ*/3, and is presumed (32) independent of 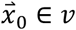 within the voxel (although it could be considered dependent on the tissue type in multicompartment models).

The RT properties of the tissue have been assumed to be constant along the trajectory of the diffusing water, thereby simplifying the above model. Given that the spatial distribution of molecules diffusing from 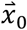 to 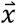 is their *initial* density, the integrals in Eq. [1] are weighted by 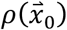. However, *w* is expected to vary in the tissue continuum along the molecule’s trajectory, which would affect the resulting signal attenuation. To account for the variations of *w*, the integral needs to be weighted by an *effective* value of *w* experienced by the molecules going from 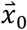 to 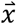 rather than by its initial value at 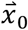. Thus, weighting the above integral by 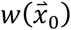, as done in the state-of-the-art dMRI models, neglects how the within-voxel variation of *w* affects the dMRI signal. Such effects could in fact be exploited to further learn about the tissue microstructure.

We propose to use an effective value for the RT weighting, *w*, to account for its change during a molecule’s diffusion. For particles going from 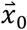 to 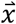 instead of weighting the integral in Eq. [1] by the initial value 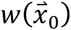, we will use the *midpoint* value,

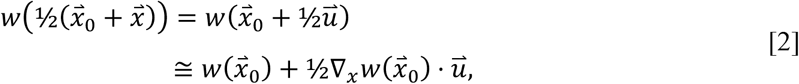

where ∇_*x*_ is the spatial gradient. This leads to:

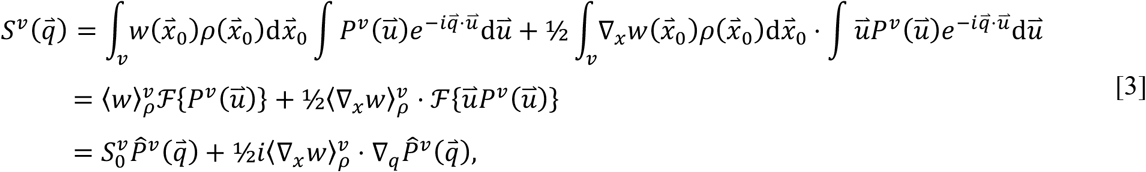

where we used the relationship 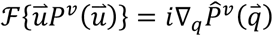, with ∇_*q*_ the gradient with respect to 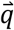 Note that our linear approximation of *w* in Eq. [2] is local only within the molecule’s trajectory and does not extend to the entire voxel.

Provided the measurements of the diffusion signal 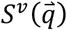 for many *q*-vectors, as is common in dMRI, Eq. [3] allows the estimation of 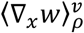, i.e. the mean ∇_*x*_*w* (weighted by PD) within the voxel, which can reveal potentially new information about the tissue microarchitecture. By contributing to the imaginary part of the signal, ∇_*x*_*w* affects both the magnitude and the phase of the signal. It has been shown that the background phase varies strongly from one diffusion image to another in the scan (due to physiological factors) (33,34), which would render the estimation of the contribution of 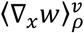 to the phase image impractical. The *magnitude* of the signal, which, in contrast, remains largely consistent from shot to shot, would be:

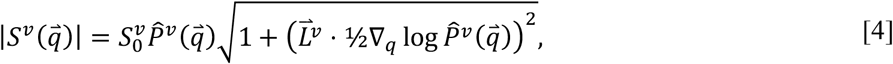

Where 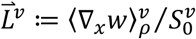, and we made the standard assumption that diffusion is symmetric (thus 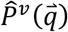 is real), as well as used 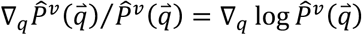. In the simple case of Gaussian diffusion, the diffusion tensor imaging (DTI) model (11) predicts 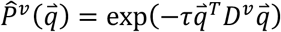, where *D*^*v*^ is the symmetric diffusion tensor at voxel *v*, resulting in the factor 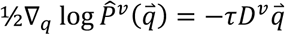 being a linear function of 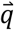 The DTI approximation, therefore, leads to:

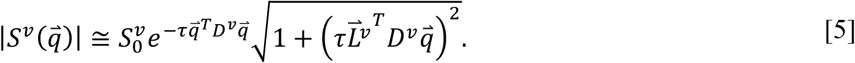

With this model, measurements of 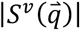 for at least 9 values of 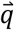 in different orientations (in addition to an 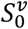 image) would be required to estimate the 6 parameters of *D*^*v*^ and the 3 elements of the vector 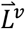 (and subsequently 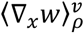). Note that, since 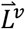 is raised to the power of 2 in Eq. [5], its sign cannot be directly recovered from 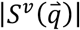 measurements, meaning that its magnitude and *orientation* (rather than direction) can be estimated. Including the fourth-order terms in 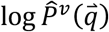 would extend the model to take advantage of diffusional kurtosis imaging (35), which is more accurate than DTI when the diffusion deviates from Gaussian (but would require 21 + 3 = 24 unknowns to be estimated from at least two q-shells).

### 2.2. Implementation and Assessment

To estimate *D*^*v*^ and 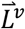 at each voxel *v*, we fit the values of the diffusion signal for all available 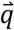 to Eq. [5] by minimizing the residual sum of squares via the pattern search algorithm (36). This nonconvex optimization could be facilitated by initializing *D*^*v*^ with standard DTI reconstruction (11), 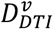, and 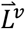 as a 3×1 vector of all zeros. However, since the objective function is symmetric with respect to 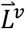 about 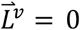, if the initial *D*^*v*^ were also the conditional minimizer of the function (with 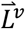 constrained to be zero, i.e. standard DTI), then the aforementioned initialization would be counterproductive, as such an initial point would already be a local optimum with an objective-function gradient of zero with respect to both *D*^*v*^ and 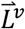. To move the initial solution away from this local optimum, we perturb the DTI-derived tensor and instead use the initial tensor 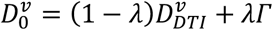, with *λ* = 10^*−*4^, where *Γ*_3×3_ is a random symmetric positive semidefinite (PSD) matrix with the same mean diffusivity as 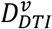. We generate *Γ* as (Wishart-distributed) *η η*^*T*^, with *η*_*3×*100_ a random matrix of independent elements drawn from the normal distribution *N* (0,1), and normalize it as 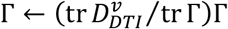. Given the convexity of the set of symmetric PSD matrices, our perturbed initial tensor, 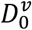, remains symmetric and PSD.

We assess the orientational accuracy of the estimated 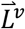 by comparing it to its discrete counterpart (used as the gold standard) computed via finite difference, testing the hypothesis that the two orientations are more aligned than expected by chance. Where the PD image, 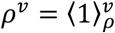, is available, the discrete counterpart of 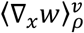 would be 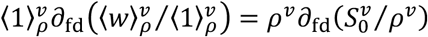, with *∂*_*fD*_ denoting the finite-difference gradient operator. The discrete counterpart of 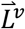 can therefore be computed as the following vector:

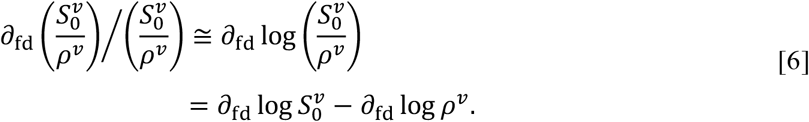

However, since the PD image is often unavailable in dMRI datasets, we approximate the above discrete counterpart with its first term, 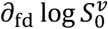. Given that the *S*_0_ and PD image intensities both correlate with the same underlying tissue, their gradient *orientations* are expected to align with each other, favoring our approximation.

We measure the acute angle (0 ≤ *θ* ≤ 90°) between the orientations of the estimated 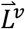 and its discrete counterpart, which should be small if the two are similarly oriented. The null hypothesis, i.e. 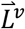 is randomly oriented with respect to its discrete counterpart, predicts the null probability density distribution of *θ* to be sin *θ* (due to the infinitesimal solid angle *D*Ω = *S*in *θ d θd ϕ*), with the mean of 57.3° (1 rad) and the median of 60°. We use two-sided *t*-tests to see if the population averages of the mean and median of 0 are different from these null values.

### 2.3. Effect Size

The diffusivity of the tissue can be quantified from 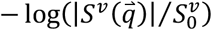. In our case, using the DTI model, that would be:

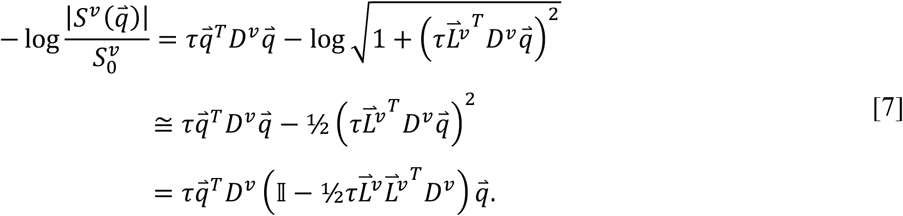

The relative contribution of 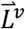 to the above quantity, which we call the dimensionless *effect size*, is of the order of 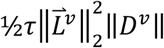. With the simplifying assumptions that PD remains relatively constant inside the voxel and *w* varies with a general linear trend, one can see 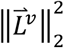 to be largely bounded by 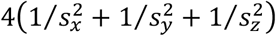, where *S* is the pixel size in the *a*-axis (these assumptions are used only here to derive this closed-form upper bound, and nowhere else). Therefore, for a dMRI of the brain white matter (WM) with *S*_*x*_ = *S*_*y*_ = *S*_*z*_ = 1.25 mm, *r* = 40 ms, and ‖*D*^*v*^‖ = 0.0007 mm^2^/*S*, the effect size would be about 10^−4^ (enabling the approximation in Eq. [7]), which might be too small to detect. This is mainly due to the small particle displacement during the diffusion time, i.e. of the order of 10 µm. The effect size can however be increased using a long *τ*. This prompted us to test our hypothesis additionally with the STE sequence (28,29) that can achieve a diffusion time of *τ* = 1 *S* (Section 2.4.2), while preserving the signal-to-noise ratio (SNR) by not increasing the effective TE (that would otherwise cause T_2_ signal loss).

It is also noteworthy that our hypothesized effect (Eq. [7]) is quadratic (rather than quartic) with respect to the q-vector, or linear (rather than quadratic) with respect to the b-value, meaning that it is not of a kurtosis (35) nature.

### 2.4. Population

We tested our algorithm on the following three datasets. Given the Gaussian-diffusion assumption in our model, for each dataset, we included only the images acquired with the lowest b-value in order to minimize signal non-Gaussianity in the data used for model estimation.

#### 2.4.1. Human Connectome Project (HCP) – Young Adult

We processed dMRI data of 617 subjects from the public WashU-UMN HCP Young Adult (27). The images had been acquired on a customized Siemens 3T scanner at the 1.25 mm isotropic resolution, with *TR*/*TE* =5520/89.5 ms and the diffusion time of *r*_*HCP*_= 40 ms, along 270 diffusion gradient directions with b-values ranging from 990 to 3010 s/mm^2^ (of which we only used b ∼ 1000 s/mm^2^), in addition to 18 b=0 s/mm^2^ (*S*_0_) images. We used the brain (gray and white matter) mask computed from the T_1_ image via FreeSurfer (37), resampled it in the dMRI space, and refined it as described in Section 2.5.

#### 2.4.2. Acquired Stimulated-Echo (STE) dMRI

We used an STE dMRI (28,29) research sequence with the long diffusion time of *τ*_*STE*_ = 1 *S* to acquire the following images on a 3T scanner (MAGNETOM Skyra, Siemens Healthineers AG, Forchheim, Germany) with a 32-channel head coil, GRAPPA acceleration 2, b-value of 1000 s/mm^2^, and isotropic voxel size of 2 mm.

We scanned 10 healthy volunteers (7 females, age: 36 ± 14 years old), whose consent and data were collected as part of an existing project to develop novel MRI acquisition software, approved by the Institutional Review Board of the Mass General Brigham. We used the receiver bandwidth of 1658 Hz/px, *TR* of 26400 ms, effective *TE* of 31 ms, and image size of 104×104×25 voxels. We collected images with 64 diffusion directions, as well as 20 repetitions of the b=0 image, resulting in a total acquisition time of 37.5 minutes. For each volunteer, as a baseline, we also acquired dMRI images with a short “effective” diffusion time of *τ*_*TRSE*_ = 22 ms, using a twice-refocused spin echo (TRSE) sequence with bipolar pulsing with *TR*/*TE* = *3*500/81 ms (and otherwise similar parameters to the STE scan), resulting in an acquisition time of 5 minutes.

In addition, we scanned an *ex vivo* sample of a brain hemisphere using the *TR* of 26200 ms, effective *TE* of 33 ms, receiver bandwidth of 1698 Hz/px, and image size of 128×128×25 voxels. To increase the SNR, we acquired images with 256 diffusion directions, with 8 repetitions per direction, as well as 320 repetitions of the b=0 image, resulting in a total acquisition time of 17.3 hours. The repeated images were averaged before further analysis.

MPRAGE T_1_ images were also acquired for all the data, from which the brain mask was extracted using FSL’s `bet` (38), transformed into the diffusion space, and further refined as described in Section 2.5. Gibbs ringing artifacts were removed from all images using MRtrix3’s `mrdegibbs` (39), and `eddy` currents and motion were corrected for using FSL’s eddy (40).

See Section 6 for the public availability of this long-diffusion-time STE dMRI dataset.

#### 2.4.3. Multi-TE (MTE) dMRI

We used the preprocessed dMRI images of the 3 subjects included in the public MTE dataset (30), acquired on a Siemens 3T Prisma scanner in 10 sessions (per subject) with 8 different *TE* values from 62 to 132 ms (with repeated measures at the shortest and longest *TE*). The dataset included two b-values (700 and 2000 s/mm^2^, of which we only used the former) with 30 gradient directions for each b-value and 4 b=0 s/mm^2^ (*S*_0_) images. The isotropic voxel size was 2.5 mm, *TR* was 5800 ms, and the diffusion time was fixed at *τ*_*MTE*_ = 20 ms. Preprocessed images had been denoised, corrected for B0 inhomogeneity, eddy current and motion correction, and aligned to the images and b-vectors to the first *TE* session. We additionally removed Gibbs ringing artifacts using MRtrix3’s `mrdegibbs` (39). We used the brain masks included in the dataset and refined them similarly to HCP, as described in Section 2.5.

### 2.5. Fiber Continuity: A Confounding Factor

Stemming from dMRI physics, our hypothesis is a relationship between diffusional information in the signal and spatial derivative of the image. However, characteristics of the fibrous tissue can additionally cause the fiber orientation to be confoundingly related to edges in the image. In particular, due to fiber continuity (41-43), fiber bundles vary smoothly along their own orientations, meaning that image edges would likely not be perpendicular to orientations with high diffusion. This means that the spatial gradient of the image in the fiber bundles is likely stronger along the lower-rather than higher-diffusion orientations.

In designing our experiments, we attempted to heuristically avoid the areas affected by fiber continuity by limiting our analysis to WM regions that are far from the bundle edges. For our *in vivo* (HCP, STE, and MTE) images, we only kept voxels with fractional anisotropy (FA) greater than 0.4, b=0 image intensity greater than 80, and mean ADC greater than 0.0002 mm^2^/s and smaller than 0.0008 mm^2^/s (HCP) or 0.0009 mm^2^/s (STE and MTE). We used a single average mask for various-*TE* images of each MTE subject. For our *ex vivo* STE image, we only kept voxels with FA greater than 0.25, mean ADC smaller than 0.0003 mm^2^/s, and b=0 image intensity greater than 90.

Furthermore, for comparison, in addition to estimating the orientation of the spatial gradient of the image using our hypothesized relationship (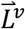 in Eq. [5]), we also estimated it following the fiber continuity assumption as the orientation of the eigenvector corresponding to the smallest eigenvalue of the diffusion tensor, calling it 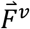.

## 3. Results

### 3.1. HCP Results

We estimated the proposed 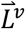, the fiber-continuity-derived 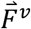, and their gold-standard discrete counterpart for each subject. Figure 1 shows the average of each of the three (normalized) vector fields across all 617 subjects, color-coded with respect to the estimated orientation. One can see that the spatial-gradient orientations derived from the two diffusion-profile-based approaches – ours (top, left) and fiber-continuity (top, right) – correspond to some extent to the discretely computed orientations (bottom, left). Furthermore, despite their similarities, there are clear regions (bottom, right) where 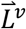 is more accurate than 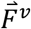 (red) and vice versa (blue).

**Figure 1.**
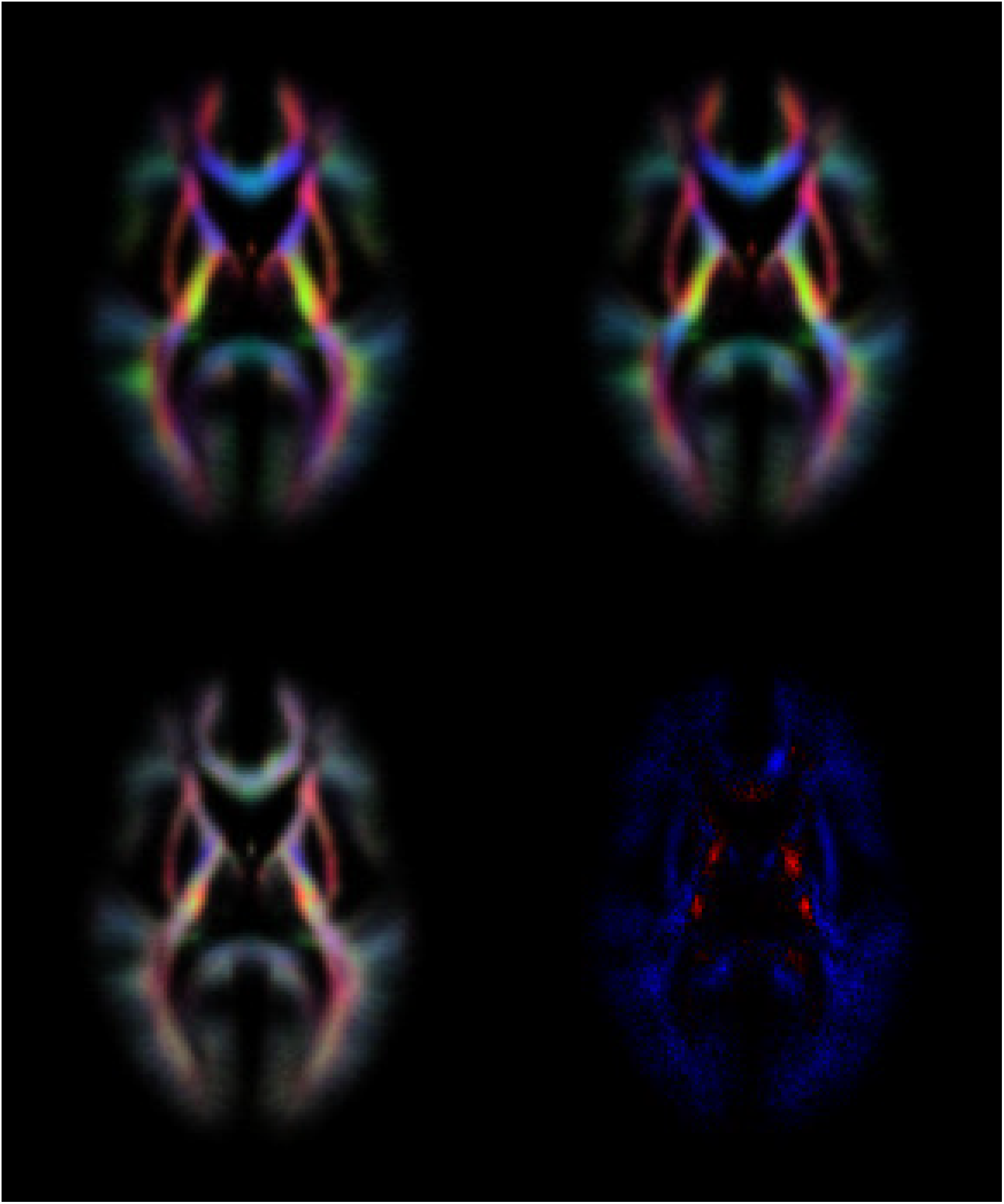
Axial view of the spatial gradient of the image estimated from the diffusion profile using the proposed (top, left) and fiber-continuity (top, right) approaches, and from the b=0 image using the finite-difference approach (bottom, left). Colors show the strength of the gradient at each spatial coordinate after normalizing the gradient, taking its absolute value, and averaging across all HCP subjects (red = R/L, green = A/P, blue = S/I; not to be confused with color-coded FA). Cross-subject average of *θ*_*L*_ − *θ*_*F*_ (bottom, right) shows regions where the proposed gradient is closer to the finite-difference one than fiber continuity is (red) and vice versa (blue).

The histogram of the acute angle *θ*_*L*_ between the estimated 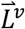 and its discrete gold-standard counterpart across all WM voxels of all subjects is plotted in Figure 2. The distribution of 0_*L*_ (blue), which is visibly shifted to the left compared to the null hypothesis (dashed curve), has a mean/median of 51.0°/51.6°, i.e. smaller than chance (57.3°/60°). A similar histogram of *θ*_*F*_ between the fiber-continuity-derived 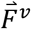 and the discrete counterpart is shown in red, which is further to the left, with its mean/median being 47.7°/47.2°.

**Figure 2.**
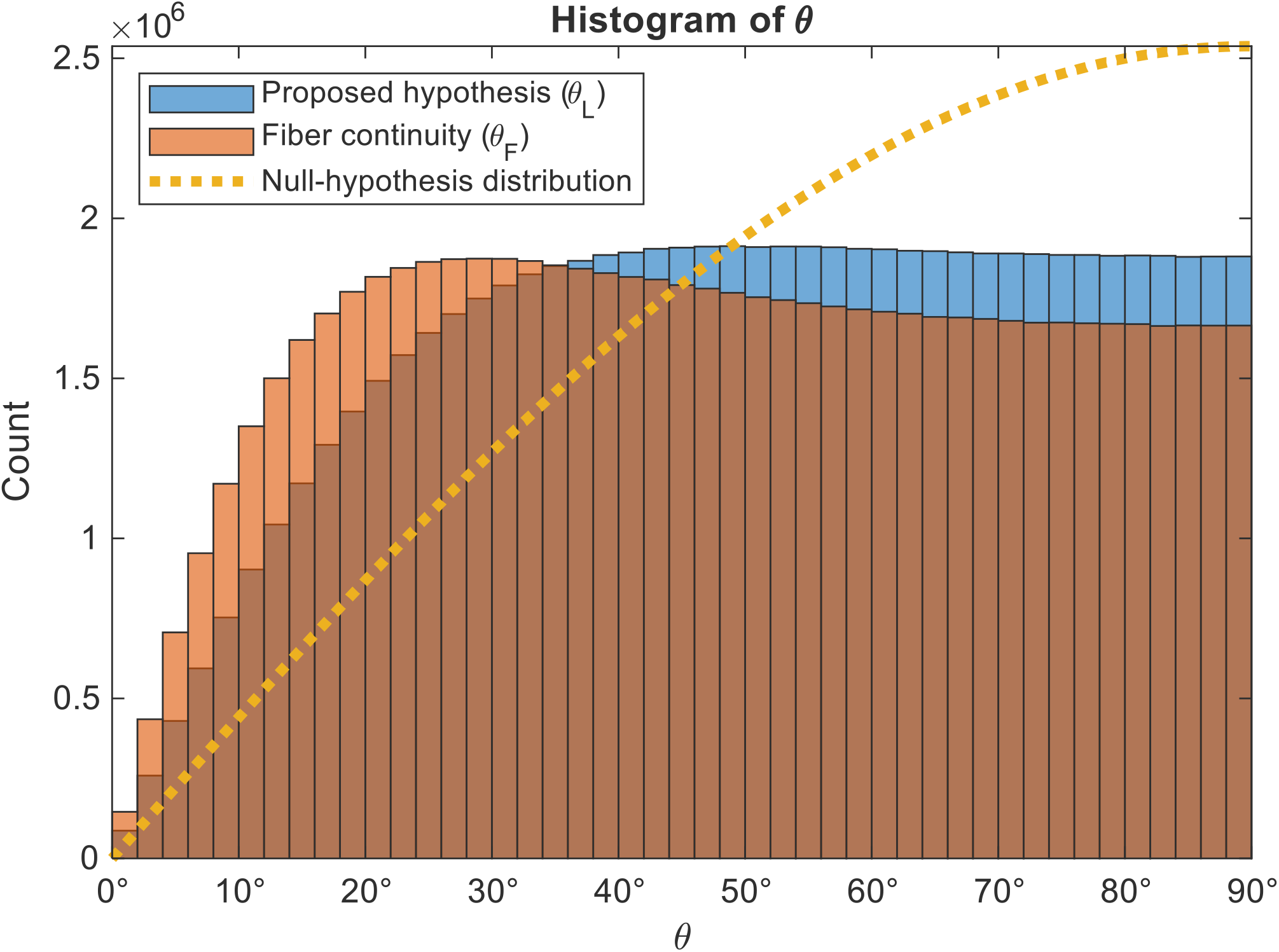
Histograms of the acute angles between spatial gradient of the image computed discretely from the b=0 image and the ones estimated from the diffusion profile using the proposed (*θ*_*L*_, blue) and the fiber-continuity (*θ*_*F*_, red) approaches, across all WM voxels of all HCP subjects. The dotted orange line is the distribution of this angle under the null hypothesis (i.e., if the two orientations were not related).

Next, we computed the mean/median of both *θ*_*L*_ and *θ*_*F*_ for each subject separately, the histograms of which across subjects are plotted in Figure 3. Two-tailed *t*-tests revealed these values to be significantly smaller than those predicted by chance (*p* = 0, within the double-precision limits).

**Figure 3.**
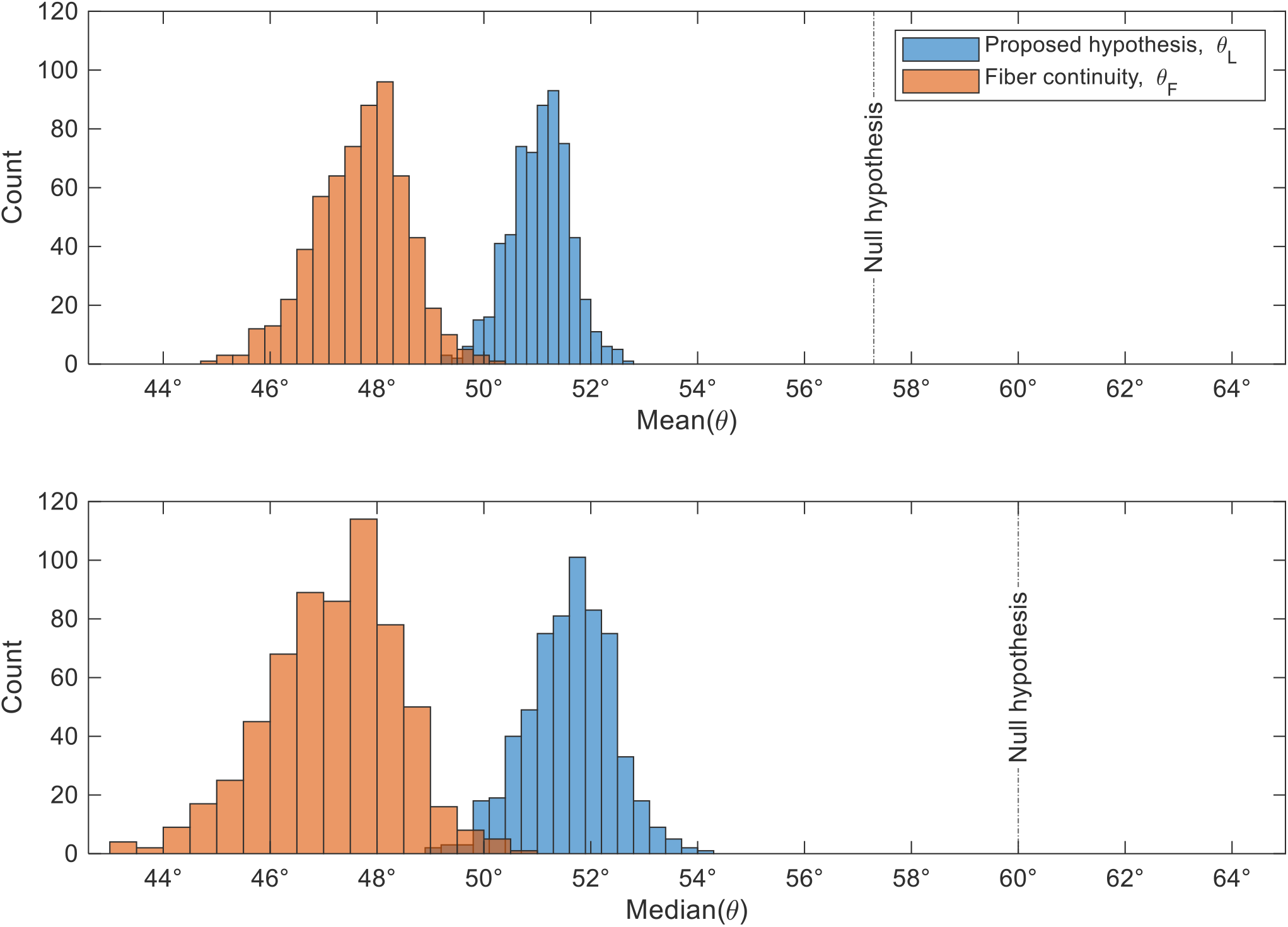
Histograms of the within-subject mean (top) and median (bottom) of the acute angles between spatial gradient of the image computed discretely from the b=0 image and the ones estimated from the diffusion profile using the proposed (*θ*_*L*_, blue) and the fiber-continuity (*θ*_*F*_, red) approaches, across all HCP subjects. The dashed vertical lines indicate the values under the null hypothesis (i.e., if the two orientations were not related).

We then compared *θ*_*L*_ and *θ*_*F*_ voxel-wise. Even though *θ*_*L*_ was on average slightly larger than *θ*_*F*_ (see above), the histogram of *θ*_*L*_ − *θ*_*F*_, plotted in Figure 4, revealed a considerable portion (44%) of voxels with *θ*_*L*_ < *θ*_*F*_, implying *distinct* orientational information mined by the two approaches (see also Figure 1, bottom, right). The small effect size when comparing the two angles is further reflected in their pairwise Cohen’s *d* = 0.16, emphasizing that, despite their population-average similarities (Figure 1), the two approaches to spatial gradient estimation are far from fully overlapping.

**Figure 4.**
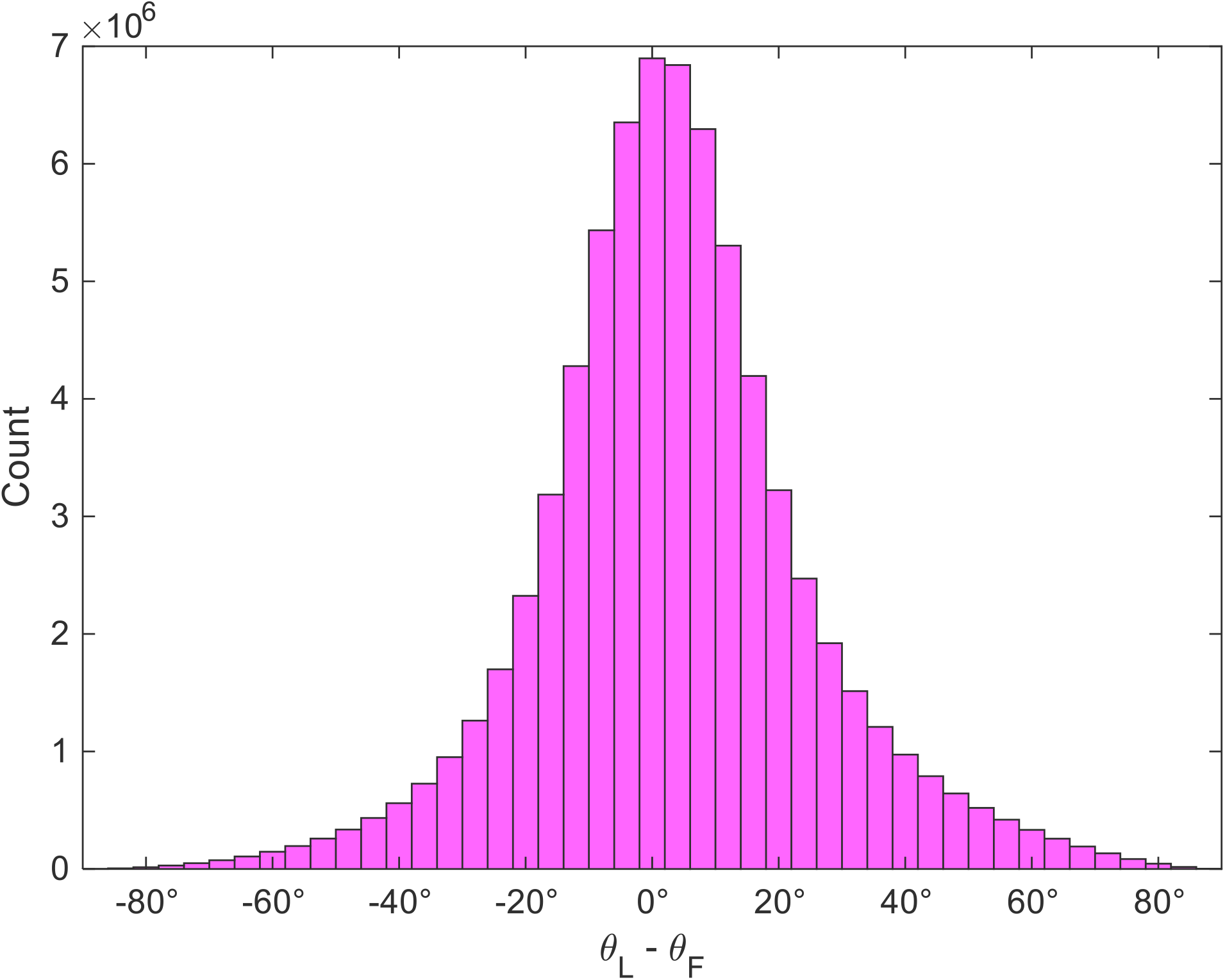
Histogram of *θ*_*L*_ − *θ*_*F*_ across all WM voxels of all HCP subjects. 44% of the voxels are on the negative side (*θ*_*L*_ < *θ*_*F*_) of the distribution.

The diffusion tensor estimated from Eq. [5] on average had 0.0087% (±0.0003%) larger Frobenius norm, 0.0104% (±0.0003%) larger mean diffusivity (MD), and 0.0071% (±0.0001%) smaller FA, compared to the tensor estimated the conventional way, i.e., by fixing 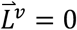 (errors are cross-subject standard error of the mean of the within-image mean value). These results are consistent with the effect size of 0.01% theoretically predicted for the HCP in Section 2.3 (but also similar to the perturbation size used in Section 2.2 to initialize the optimization).

### 3.2. STE Results

Figure 5 shows the q-ball orientation distribution functions in constant solid angle (CSA-ODFs) (44) reconstructed with a (real and symmetric) spherical harmonic order of 4 and visualized using our public MATLAB (The MathWorks Inc., Natick, MA, USA) toolbox (45) (see Section 6) for a representative subject (the one with median θ_*L*_) in our *in vivo* STE dataset.

**Figure 5.**
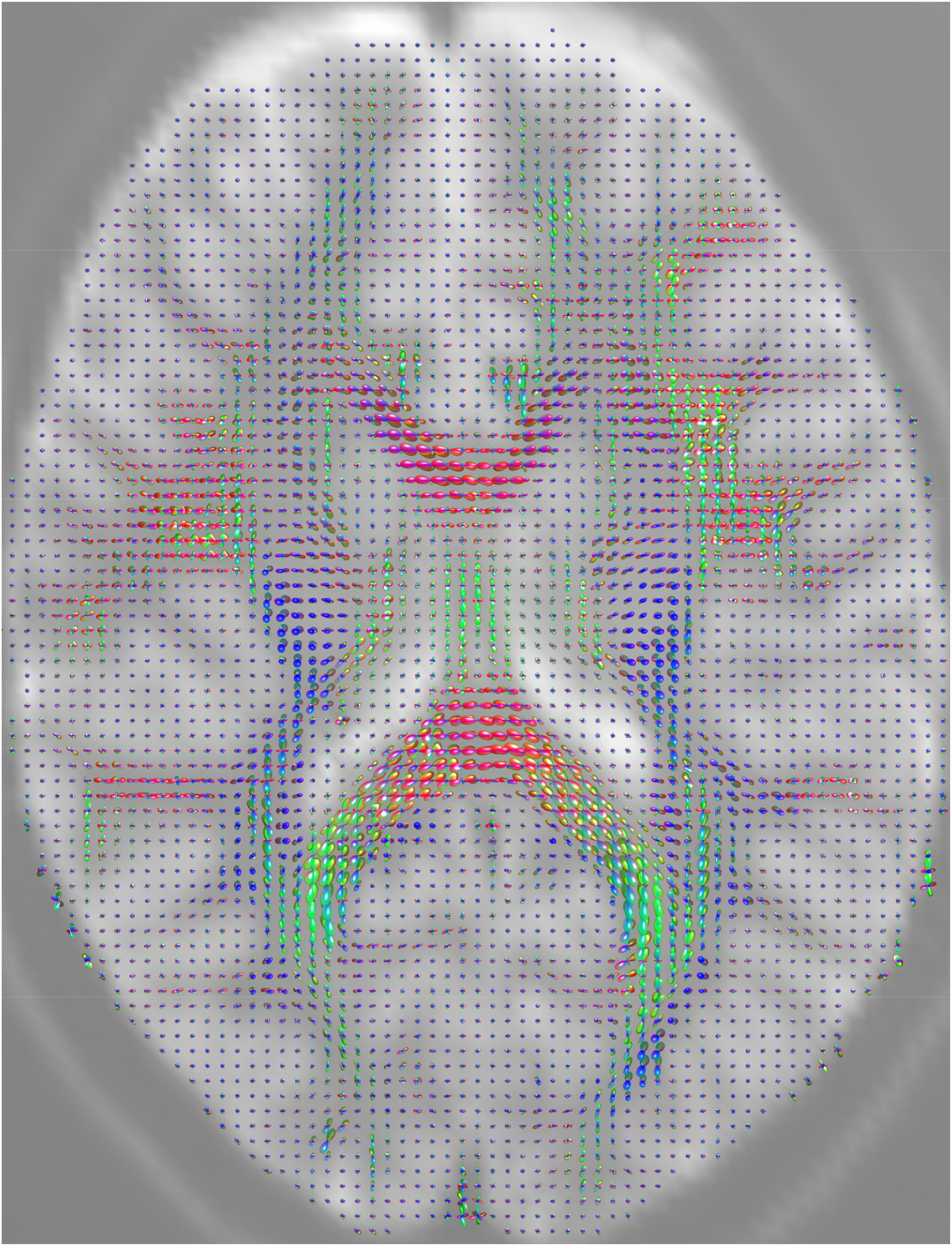
Axial view of the q-ball CSA-ODFs reconstructed from the STE data of a representative participant.

An analysis similar to Section 3.1 for our *in vivo* STE images revealed the mean/median of *θ*_*L*_ and *θ*_*F*_ to be 47.4°/46.7° and 43.3°/40.8°, respectively, which are again smaller than chance (57.3°/60°). Within-subject mean/median of both *θ*_*L*_ and *θ*_*F*_ were significantly smaller than those predicted by chance (cross-subject *t*-test *p* < 10^-12^ and *p* < 10^-13^, respectively). In contrast, the experiment on our short-*τ* TRSE dMRI images resulted in the mean/median of *θ*_*L*_ and *θ*_*F*_ to be 50.8°/51.4° and 48.0°/47.6°, respectively, i.e. larger than the STE values but smaller than chance. Within-subject mean/median of both *θ*_*L*_ and *θ*_*F*_ were larger than those of the STE for every subject, but overall significantly smaller than chance (*p* < 10^-9^). Comparing 0_*L*_ and 0_*F*_ voxel-wise revealed 42% and 45% of voxels with *θ*_*L*_ < *θ*_*F*_ in the STE and TRSE experiments, respectively.

The diffusion tensor estimated from Eq. [5] for the STE and TRSE images, respectively, on average had 0.07% (±0.02%) and 0.005% (±0.002%) larger Frobenius norm, 0.08% (±0.02%) and 0.006% (±0.002%) larger MD, and 0.038% (±0.004%) and 0.0046% (±0.0003%) smaller FA, compared to the tensor estimated the conventional way. These are roughly consistent with the theoretically predicted effect sizes of 0.1% and 0.002% for the STE and TRSE data, respectively (Section 2.3). To ensure that this difference was not simply due to the different *τ* values that we used in Eq. [5] during the pattern search optimization for the two experiments, we refit the TRSE data, but this time using *τ* = 1 s in the optimization, which still resulted in an effect size 7 times smaller than that of the STE (and no change in 0_*L*_).

Regarding our *ex vivo* STE image, the CSA-ODFs reconstructed from the diffusion tensors are depicted in Figure 6. The mean/median of θ_*L*_ and θ_*F*_ for this image were 54.6°/56.6° and 52.9°/54.4°, respectively, which are smaller than chance (but not by as much as the *in vivo* results), with 47% of voxels showing θ_*L*_ < θ_*F*_. The diffusion tensor estimated from Eq. [5] on average had 0.001% larger Frobenius norm, 0.001% larger MD, and 0.003% smaller FA, compared to the conventional tensor. These values are much smaller than the case of *in vivo* STE, possibly due to the smaller water diffusivity in the *ex vivo* tissue and lower SNR.

**Figure 6.**
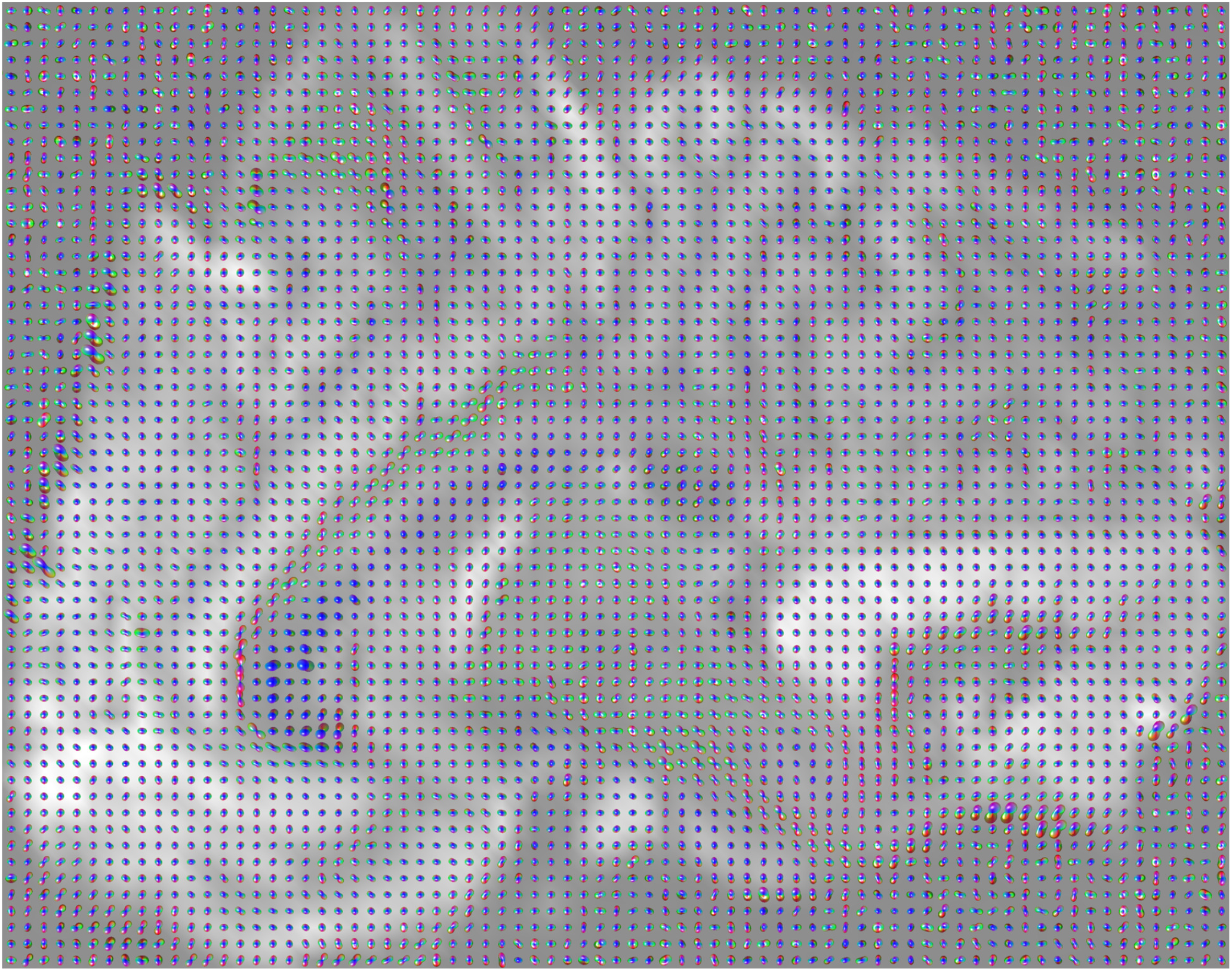
Sagittal view of the DTI CSA-ODFs reconstructed from the *ex vivo* STE scan.

### 3.3. MTE Results

An analysis similar to Section 3.1 over all subjects resulted in mean/median of θ_*L*_ and θ_*F*_ of 50.3°/50.7° and 47.6°/47.0°, respectively, i.e., smaller than chance. Comparing θ_*L*_ and θ_*F*_ voxel-wise revealed 45% of voxels with *θ*_*L*_ < *θ*_*F*_.

Given the long *TR* of the data, the RT weighting would chiefly reflect the T_2_ weighting, 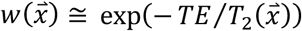, resulting in 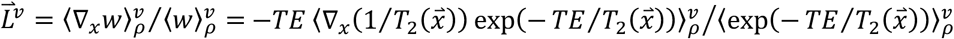, with the magnitude 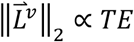 (roughly). We indeed observed that the within-image median of our estimated 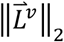 generally increased with *TE*, as shown in Figure 7. A linear mixed-effects analysis, relating median 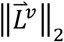 to *TE*, with the subject as the random effect, revealed a significant positive relationship (*p* = 10^-7^). The plot, however, shows some quadratic behavior especially at low *TE*, which might be due to other ways that varying the *TE* affects the signal. For instance, our single-compartment approximation may give rise to nonlinear *TE* dependencies in the presence of multiple compartments, particularly at low *TE* where short-T_2_ compartments become more influential. We also observed that MD monotonically decreased here with *TE* on average from 0.000753 to 0.000716 (±0.000003) mm^2^/s.

**Figure 7.**
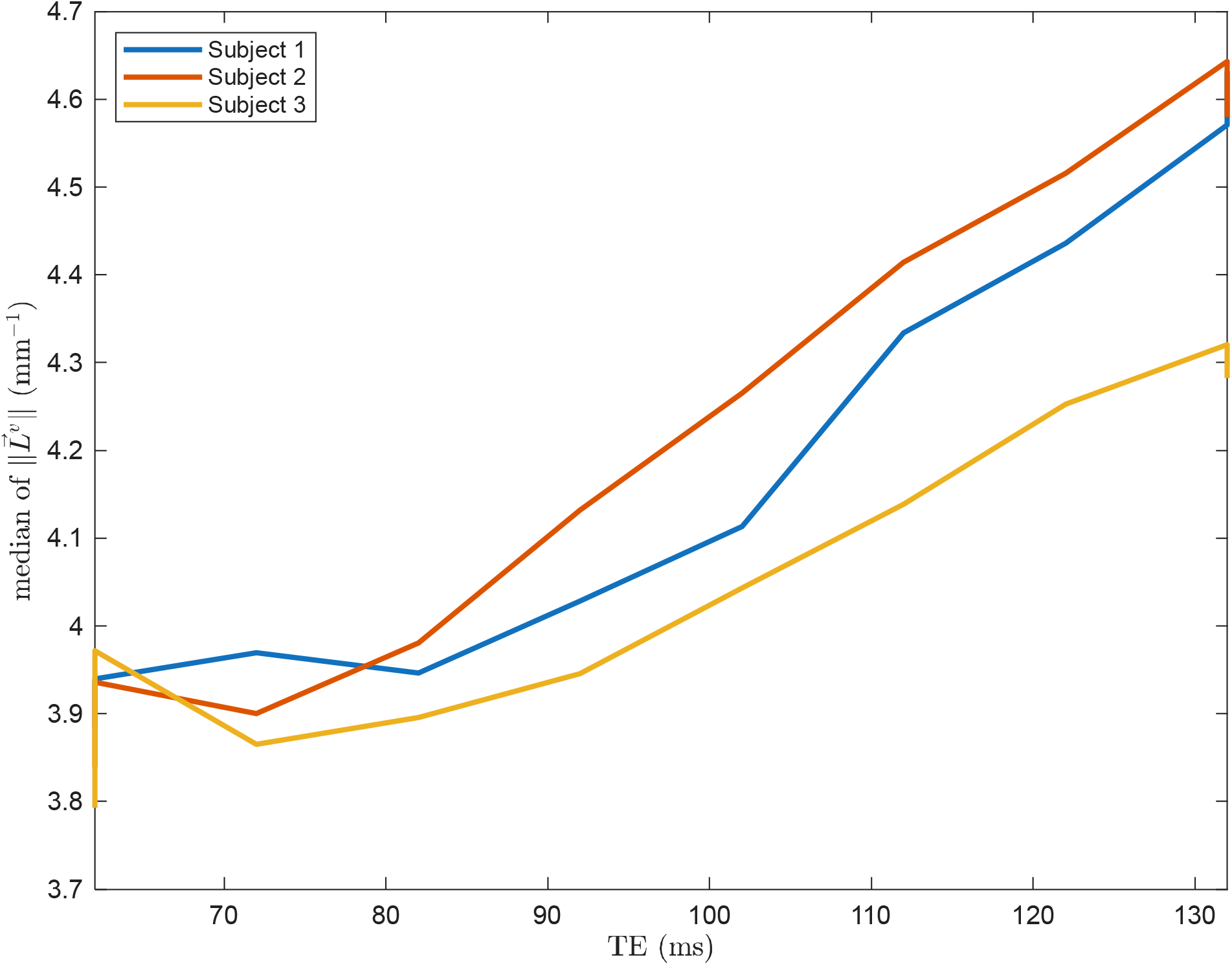
Within-image median of the magnitude of the estimated 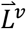 in the MTE dataset for each *TE*.

The diffusion tensor estimated from Eq. [5] on average had 0.00134% (±0.00004%) larger Frobenius norm, 0.00224% (±0.00006%) larger MD, and 0.009% (±0.006%) smaller FA, compared to the conventionally estimated tensor (median value across different-*TE* images was used for each subject). These are consistent with the theoretically predicted effect size of 0.001% (Section 2.3).

To compute the finite-element gradient in the above, we used the average *S*_0_ image across all (various- *TE*) images of each subject. Next, we applied MATLAB’s exponential curve fitting tool to instead fit those images to 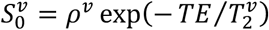 to estimate 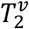 and the PD, *ρ*^*v*^, for each voxel *v*, with positivity constraints for both variables. Unfortunately, the resulting 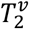 (and 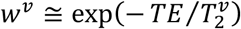) maps were not of great quality, as the estimated *w*^*v*^ had slice-wise intensity inhomogeneities that compromised the accuracy of our finite-difference estimate of the gradient of *w*^*v*^, i.e. *∂*_*fD*_*w*^*v*^, at least in the slice-select direction. We computed the within-image median of *θ*_*L*_ for each subject and *TE*, but this time using the estimated *w*^*v*^ or *ρ*^*v*^ as the gold standard. For one subject, this did not improve (decrease) *θ*_*L*_ over the unseparated 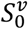, whereas for the other two subjects, the PD image resulted in a slightly (∼0.3°) decreased *θ*_*L*_.

## 4. Discussion

The spatial gradient of the RT weighting stems from local variations in the molecular composition of the biological tissue (water, fat, macromolecule, and paramagnetic substance contents), which are generally due to the spatial variations in the biophysical properties of the healthy tissue, but can also be a result of heterogeneities caused by pathology (edema, tumors, hemorrhage, demyelination, etc.) (15,46-48). We have introduced a model that takes advantage of diffusional information in the dMRI signal to infer new spatial information, thereby deriving a new relationship between the diffusion profile and the spatial gradient of tissue RT weighting. We evaluated our model on 617 brain dMRI images from the public HCP database, on 10 *in vivo* and 1 *ex vivo* brain STE images that we acquired at our Center, and on 30 brain dMRI images (with various *TE*s) of 3 subjects from the MTE dataset. We observed the effect of our hypothesized relationship in all our results; namely, the spatial-gradient orientation estimated from the diffusion profile using our relationship was statistically significantly closer to the gold-standard orientation (approximated through finite difference) than predicted by chance.

While the relationship between diffusional and RT properties of the tissue has been discussed (12-15) and measured (16-21,49) in the literature, our proposed framework is – to the best of our knowledge – the first to enable the estimation of the within-voxel spatial gradient of the RT weighting from the dMRI signal. In the work most relevant to ours (15), the authors present a detailed mathematical framework that predicts how the measured dMRI signal is affected by the diffusion time, the intrinsic variations of the Larmor frequency, and the intrinsic variations of the transverse relaxation rate, 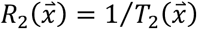. For the latter case, they derive a general integral formula containing the Fourier transform of the autocorrelation function of 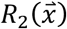. Our mathematical approach, however, is distinct from that of (15); in particular, we calculate the effects of variations of the entire RT weighting, 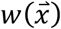, thereby naturally accounting for not only the transverse 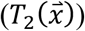 but – as opposed to (15) – also the longitudinal 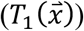 RTs (although the effects of the latter are small in dMRI). Furthermore, we derive a closed-form formula (Eq. [5]) that enables the direct estimation of the spatial gradient of 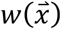, whereas, to the best of our knowledge, the derivations in (15) do not provide a straightforward means to estimate the spatial gradient (of 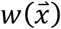 or 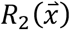).

The decision to test our hypothesis additionally on the STE images was motivated by the theoretical prediction that the spatial image gradient affects the signal more strongly at longer diffusion times (Section 2.3). Our STE protocol with *τ*_*STE*_ = 1 *S* provided us with a diffusion time 45, 25, and 50 times longer than those of our TRSE (*τ*_*TRSE*_ = 22 ms), the HCP (*τ*_*HCP*_ = 40 ms), and the MTE (*τ*_M*TE*_ = 20 ms) images. The predicted gradient orientation was indeed more accurate in the case of the STE images than in all other images, potentially due to the longer diffusion time. Importantly, the relative changes in the diffusion metrics stemming from the spatial gradient – while possibly overall influenced by the amount of perturbation in the initial tensor, i.e. the dataset-independent *λ* in Section 2.2 – varied for different datasets consistently with our theoretical predictions.

A confounding factor in validating our hypothesis was the concept of fiber continuity (41-43), a characteristic of the fibrous tissue that implies smooth variation of a fiber bundle along its orientation, hence smaller image gradient along high-diffusion orientations (see Section 2.5). To distinguish the relationship derived from our dMRI-physics hypothesis from that due to fiber continuity, we also estimated the spatial gradient of the image from the latter relationship, which produced, on average, more accurate spatial-gradient orientations (when compared to the gold standard) than our hypothesis did. Nevertheless, we observed that in a substantial portion of the voxels (44% in HCP, 42% in *in vivo* STE, 47% in *ex vivo* STE, 45% in TRSE, and 45% in MTE), our hypothesized relationship still led to more accurate orientation estimation than fiber continuity did (an example is depicted in Figure 1, bottom, right). This implies that, regardless of the fiber continuity effect being stronger than our hypothesized effect, the latter is not simply a weaker version of the former. In other words, our observations rule out the (full) overlap of the two effects, which can be clearly observed in the broad distribution of the difference between the acute angles estimated from our and fiber-continuity approaches (Figure 4).

The diffusion tensor estimated from our model showed nominally higher diffusivity compared to the conventional tensor. This is expected, since the square root in Eq. [5] is greater than 1, and a higher diffusivity in the exponential is needed to compensate for that. Our tensors were also slightly more isotropic, which could be because some of the anisotropy in the signal was accounted for by the new factor related to the spatial gradient. Nevertheless, these differences were minute (<0.01% in conventional dMRI and <0.1% in STE) and well below the noise level of standard DTI estimation. This implies that the impact of accounting for the spatial gradient (i.e., including the square root factor in Eq. [5]) on the estimated diffusion tensors and derived metrics is, as predicted in Section 2.3, minimal.

Potential future directions would be to investigate whether the spatial information hidden within the diffusion profile could: 1) help to improve the characterization of fine WM pathways and cortical microstructure, especially given that the spatial gradient for a voxel is estimated at the microscale (rather than the mesoscale as the finite-element approach does using neighboring voxels), 2) be used as a dMRI biomarker for disease, and 3) be exploited as supplemental input to improve dMRI super-resolution algorithms (50-53), given that our microscopic and the finite-element mesoscopic approaches estimate the gradient using distinct sources of information, thereby complementing each other in jointly informing the super-resolution method about the image spatial gradient. The latter is especially important as higher dMRI spatial resolution reduces the unwanted mixture of tissues in a voxel (partial volume effect), enhancing the precision and reliability of diffusion modeling and tractography (7,54-56).

Our STE dMRI dataset, which we make publicly available (see Section 6), is unique in that it has been acquired with an unusually long diffusion time of *τ*_*STE*_ = 1 s. This dataset includes *in vivo* brain images of 10 healthy volunteers and 1 *ex vivo* brain image. While we collected these data specifically to test the hypothesis proposed here, we hope that this dataset will be useful for the research community to study the diffusion properties of the neural tissue at very long diffusion times (57), e.g. the restricted diffusion (58), and test other hypotheses. Public STE datasets by other groups, e.g. of the human heart (59) or the mouse spinal cord (60), can further facilitate such efforts.

### 4.1. Limitations

Although we showed that the effects of the proposed model did not fully overlap with those of fiber continuity, our significant results may still be due to a mix of the two effects. Further evaluation, e.g. on physical phantoms constructed while minimizing the fiber continuity effect, can be helpful to conclusively validate our hypothesis.

In our evaluation, due to the lack of the “ground-truth” spatial gradient, we resorted to a surrogate gold standard, i.e. the finite-difference gradient, which, although informative, is not perfect. We further made an approximation by ignoring the variations in the PD image in Eq. [6]. Nonetheless, since we used only the orientation (but not the magnitude) of the finite-difference gradient for validation, and given that the gradients of the PD and b=0 images are both related to the same underlying tissue and thus oriented similarly to each other, the aforementioned approximation is not expected to have substantially affected the accuracy of our validation. Another consideration about our evaluation is that the gold-standard gradient was computed as the *central* finite difference, which, by using multiple voxels, introduces some blurring that is not expected in our diffusion-based model, hence a discrepancy in the comparison. While our results show a statistically significant relation between the gradients estimated through our and the finite-element methods, the effect size was not large *per se* (as predicted in Section 2.3). Our approach in fact quantifies the spatial gradient using within-voxel microscopic information hidden in the diffusion signal, whereas the finite-element approach does so using mesoscopic information from neighboring voxels. Therefore, the two measures are of different natures and affected by distinct sources of error.

To test our hypothesis, we applied the DTI model due to its simplicity and widespread use. This model’s ability to represent the diffusion signal, however, is limited in regions with fiber crossing and where the diffusion profile deviates from Gaussian (especially at high b-values). We attempted to alleviate these by limiting our analysis to the lowest b-value in each dataset, as well as to regions with FA values above a certain threshold. Nevertheless, extending Eq. [5] to higher-order models, such as diffusional kurtosis imaging (35), can further help to overcome the limitations of DTI. Systematic effects such as B1 field inhomogeneity and point spread function may also have affected the gradient estimated via both our and the finite-element approaches, which should be considered when interpreting our results.

When comparing the STE to standard dMRI, one must consider the lower SNR (two-time reduction in signal strength) inherent to STE (29) and the inadequacy of the DTI (Gaussian) model at long *τ* (61), both of which could have potentially affected our results. Furthermore, in our subject-wise comparison of the STE and TRSE images, the potential confounding factor of varying eddy current distortion levels may have put our STE images at an advantage (especially since even fiber-continuity results were better for STE). Lower eddy current distortion is in fact expected in our STE sequence due to its weaker diffusion gradients (compensating for the longer diffusion time to maintain the same b-value). This concern about the comparison is, however, alleviated thanks to the TRSE sequence with bipolar pulsing, which is effective in mitigating eddy currents (62), as well as our additional post-acquisition correction for such distortions. Another limitation of our comparison is the existence of some residual concomitant field in our bipolar TRSE images (63).

## 5. Conclusions

We extended the standard dMRI reconstruction model to account for within-voxel variation of the RT properties of the tissue, and derived a closed-form relationship in the case of DTI. The new mathematical model enables the estimation of the spatial image gradient from the diffusion profile. We evaluated our model on the public HCP and MTE datasets, as well as a unique in-house STE dataset that we acquired with a very long diffusion time (and provide to the public). Our experimental results support the validity of our hypothesis with statistical significance. Future work consists of leveraging the estimated spatial gradient to learn about tissue microstructure, discover biomarkers, and increase the spatial resolution of dMRI.

## 6. Data and Code Availability

Our anonymized STE dataset, described in Section 2.4.2, will soon be publicly available at: www.nitrc.org/projects/ste Our MATLAB code for the proposed spatial-gradient estimation is included as the `estimateSpatialGradient` function in our public CSA-ODF and Hough-Tractography toolbox (45) (www.nitrc.org/projects/csaodf-hough). Diffusion and STE fiber orientation reconstruction was performed using the same toolbox.

The public WashU-UMN Human Connectome Project (HCP) Young Adult (27) database is available at: https://www.humanconnectome.org/study/hcp-young-adult/data-releases The public Multi-TE (MTE) dataset (30) is available at: https://doi.org/10.57760/sciencedb.o00133.00038

## Acknowledgment

We would like to thank Matthew Vera for his help in acquiring the *ex vivo* brain image. Support for this research was provided by the Michael J. Fox Foundation for Parkinson’s Research (MRI Biomarkers Program award MJFF-021226) and the National Institutes of Health (NIH), specifically the National Institute on Aging (RF1AG068261, R01AG068261). Additional support was provided in part by the BRAIN Initiative Cell Census Network grant U01MH117023, the National Institute for Biomedical Imaging and Bioengineering (P41EB015896, R01EB023281, R01EB006758, R21EB018907, R01EB019956, P41EB030006), the National Institute for Neurological Disorders and Stroke (U24NS135561, R01NS0525851, R21NS072652, R01NS070963, R01NS083534, U01NS086625, U24NS10059103, R01NS105820), the National Institute on Aging (R01AG079422, R01AG064027, R01AG008122, R01AG016495, R01AG070988), the National Institute of Mental Health (R01MH121885, RF1MH123195), and the NIH Blueprint for Neuroscience Research (U01MH093765), part of the multi-institutional HCP. Computational resources were provided through a Microsoft Azure Credit Grant by the Harvard Data Science Initiative, the Massachusetts Life Sciences Center, and NIH Shared Instrumentation Grants (S10RR023401, S10RR019307, S10RR023043). B. Fischl is a medical advisor to DeepHealth, a company whose medical pursuits focus on brain imaging and measurement technologies. His interests were reviewed and are managed by Massachusetts General Hospital and Mass General Brigham in accordance with their conflict-of-interest policies. T. Feiweier is employed by, owns stocks of, and holds patents filed by Siemens Healthineers AG.

